# Applying telecentric stereo 3D-measurement to small Lepidopterans – bridging the macro and the microscale with isotropic micrometer resolution

**DOI:** 10.1101/2025.05.15.654204

**Authors:** Andreas Walter Stark, Adeoluwa Osadare, Matthew Guo, Gregor Joerg Gentsch, Dennis Boettger, Gunnar Brehm, Christian Franke

## Abstract

We present a straightforward, application-driven telecentric stereo 3D-measurement system for high-precision measurements, designed for applications ranging from industrial quality control to biological research including scanning of Lepidoptera moths. Utilizing a dual-camera setup with telecentric lenses and structured illumination, our system achieves lateral resolution of 8.0 *μm* and axial resolution of 4.46 *μm* in a measurement volume of 11 *mm* × 11 *mm* × 6 *mm*. We address challenges typically encountered when using standard libraries like OpenCV, e.g. in extrinsic parameter estimation using a dedicated calibration method that corrects for a potential model mismatch due to telecentricity. Our approach adapts existing methods, such as telecentric stereo vision and structured illumination, into an optimized, user-friendly system tailored for life science research, enabling detailed 3D reconstructions of scattering objects, such as small moths, with isotropic micrometer accuracy. This work presents an application-driven approach for biological 3D metrology by integrating existing technologies (telecentric stereo vision, structured illumination) into a specialized imaging platform suitable for non-invasive morphological studies. Unlike conventional CT or microscopic approaches, our method provides a balance of precision, scalability, and practical usability for non-expert users with the aim to study developmental changes in species under varying environmental conditions, while also methodically bridging the gap between macroscopic and microscopic resolution in biological imaging.

## Introduction

Accurate three-dimensional (3D) imaging of biological specimens is essential for revealing how organisms sense, move, and adapt to their environment, be it through the fine whiskers of rodents^1,2^ or the fragile components of insects^3,4^, like wings of moths (Lepidoptera)^5,6^. These intricate structures are often soft, light-scattering, and easily deformed, requiring non-contact measurement techniques to capture their morphology without causing damage. The accurate 3D measurement of biological specimens, such as small rodents^1,2^ and insects^3^, e.g. moths (Lepidoptera)^5^, is critical for understanding how, for example, multisensory integration in mammals’ functions and also how species adapt to environmental changes. Such fine and delicate structures (whiskers, wings, etc.) should be measured with a contactless method, due to them being prone to damage or distortion. These measurements should also be conducted in a reasonable time frame, allowing for statistically relevant sample sizes. Traditional (stereo-) photogrammetric methods are typically limited to lateral resolutions above 50 *μm* (laterally)^7–10^ - although some can reach a resolution around 10 *μm*^11,12^, while microscopic techniques, though offering sub-micrometer resolution, are constrained by a small field of view - often not exceeding 2 *mm*. For instance, confocal microscopy typically provides a lateral resolutions in the range of 0.2–1.0 *μm* but is limited to volumes *<*1 *mm*^3^. Computer tomographic (CT) methods, while satisfying most conditions, are time and resource-intensive for larger studies and often lack the sensitivity to resolve subtle biological structures^13^, even though the achieved resolution can reach 10 *μm* isotropically. While there have been telecentric approaches to 3D-measurement, allowing principally a high resolution and still a macroscopic field of view^14–17^, biological applications using standard libraries and structured illumination remain underexplored in this segment of metrology. Here, we aim to lower the entry barrier for life scientists to utilize high-quality 3D measurements, opening up new avenues of quantification. In this study, we introduce a telecentric stereo 3D measurement system that overcomes these limitations. While avoiding problems of standard open-source libraries such as OpenCV^7,8^ that are designed for pinhole-based camera models, we provide an in principle sub-cellular, isotropic resolution suitable for applications in both macro- and microscopic domains. We show that telecentric stereo-imaging, in combination with structured illumination, enables precise volumetric reconstructions with isotropic single digit micrometer resolution. To achieve this, we have adapted existing methodologies and provide a streamlined measurement system for biological research, reducing the complexity of existing optical metrology techniques. Finally, we demonstrate the system’s applicability in biological questions through the 3D measurement of a small moth specimen (Geometridae: Idaea sp.), capturing subtle details that could e.g. provide insights into morphological adaptations of insect species along environmental gradients^5^. Small Lepidopterans exhibit fragile, complex wing structures and subtle morphological variations, making them an ideal test case for high-precision, non-contact volume measurement. Unlike CT, which requires extensive resources, preparation and measurement time, and confocal microscopy, which has limited field of view, our method allows quick, high-resolution scanning of macroscopical organisms with minimal setup effort. The measurement volume of 11 *mm* × 11 *mm* × 6 *mm* together with a nearly cubic isotropic resolution effectively bridges the macroscopic and the microscopic scale.

## Results

### Reconstruction of Lepidopterans

Within this work a telecentric 3D-measurement setup with active, structured illumination was realized and adapted. The measurement volume had dimensions of 11 *mm* × 11 *mm* × 6 *mm* and allowed a lateral resolution of 8 *μm* and a precision of 4.46 *μm* in axial direction. To demonstrate the applicability and sensibility of this high-precision 3D measurement method to biological samples, we measured a moth specimen (Geometridae: Idaea sp.) with the system. This specific sample is of interest due to its delicate nature, its fine details and the morphological adaptations of moths that occur as a reaction to different living conditions. An exemplary visualization of one of the reconstructions can be seen in Fig. 1.

**Figure 1.**
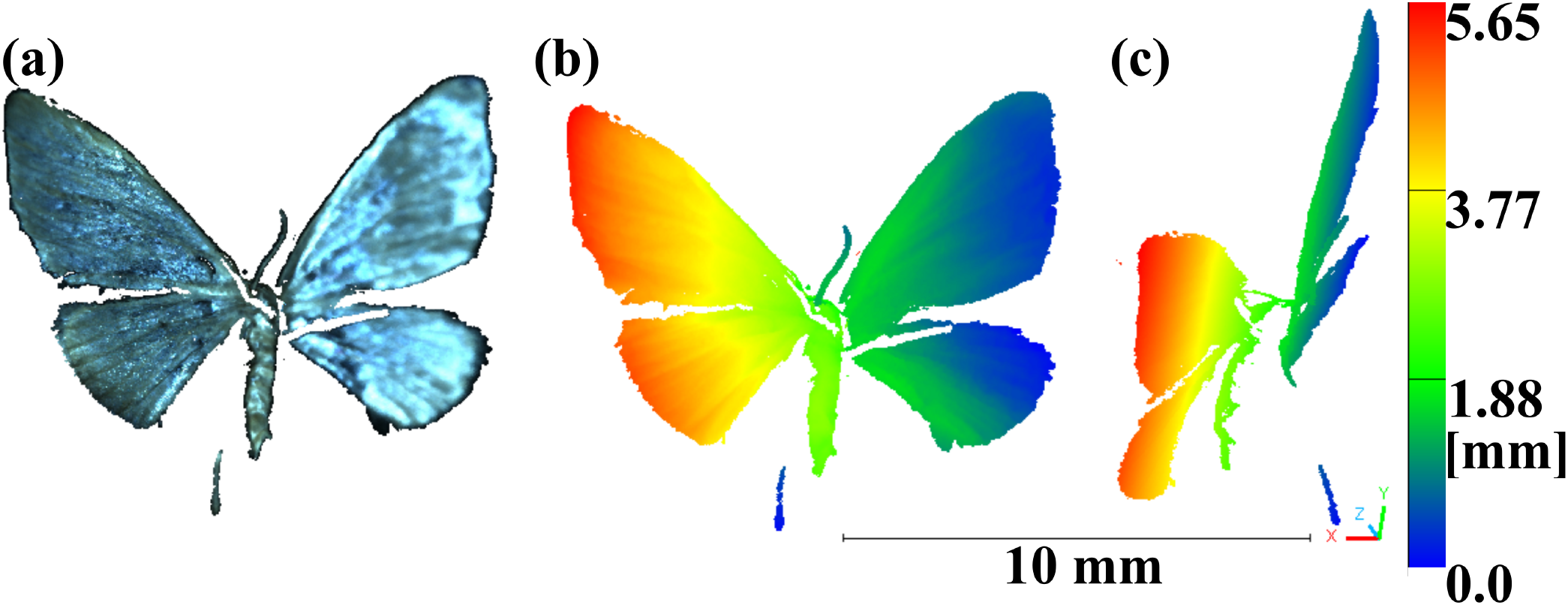
(a) 3D-reconstruction of a small moth species (Idaea sp.) in front view and real colors, (b) front view with z coordinates as color, (c) side view with z coordinates as color. Scale bar and color bar in mm. Point clouds are available in the supplementary file^18^.

Even though the tiny specimen had a length of below 4.4 *mm*, a maximum width of 0.75 *mm* and a wingspan of 10.75 *mm*, Fig. 1 showcases several distinct features of the moth, reconstructed in high detail. The measurement procedure was repeated several times for different positions, the resulting point clouds are available in the supplementary file^18^. To give an example of the advantage of this 3D-measurement techniques of small specimen as compared to measurement in 2D^4^ we look into the images and the reconstruction of an antennae of the moth (see Fig. 2). Here we show the full view of the first camera under homogenous illumination (Fig. 2) (a)) and enlarged the antennae with measurement spots in 2D, to retrieve the geometrical length as in^4^ (Fig. 2) (b)). We then used CloudCompare^19^ to fit a polyline on the reconstruction of the antennae, stemming from the 3D-measurement in this position.

**Figure 2.**
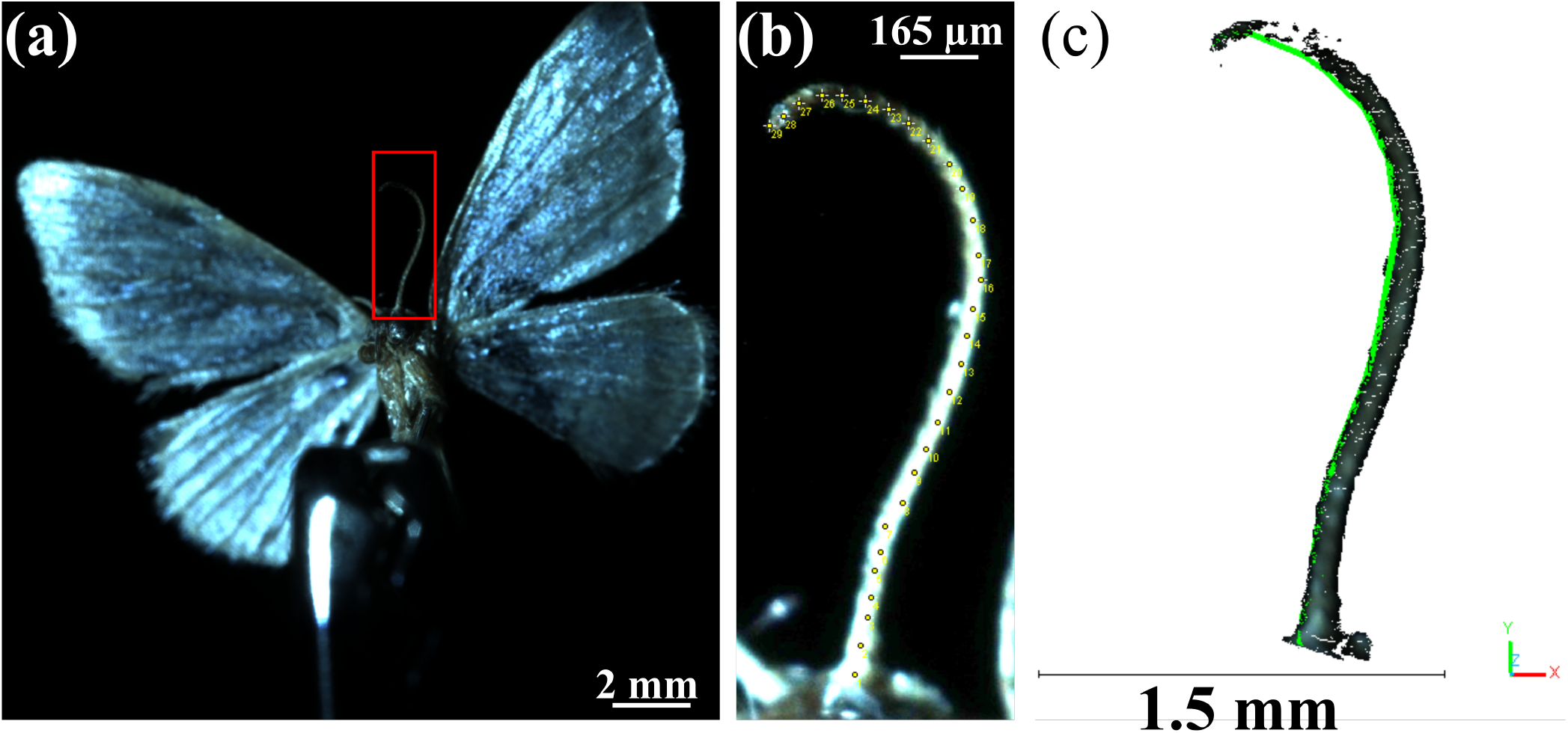
(a) Single image with homogeneous illumination from the first camera of the same moth as in Fig. 1 but under a slightly different angle. The image part containing the antennae of the moth is outlined in red. (b) Cropped image part containing the antennae increased in brightness to improve visibility. Comparable to^4^ a 2D-measurement of the length of the antennae was performed using Fiji/imageJ. The resulting length (each pixel representing 4 *μm*) was calculated to 2.576 *mm*. (c) Crop of the 3D-reconstruction of the moth under this view, containing the antennae. Using CloudCompare a polyline was fitted on the point cloud. The length of the polyline and the points representing the antennae accordingly was measured to be 3.733 *mm*. Point clouds are available in the supplementary file^18^.

A comparison between the 2D image and the 3D reconstruction of the moth’s antennae revealed a significant discrepancy in the measured length: while the 2D projection suggests a length of approximately 2.576 *mm*, the corresponding 3D polyline measures around 3.733 *mm*. This difference arises from the inherent curvature of the antennae in three-dimensional space, which is not captured in the 2D image due to the foreshortening effect introduced by the projection. Consequently, relying on 2D measurements alone would result in a substantial underestimation of true anatomical dimensions. This point cloud data can enable the detailed quantification of features at the micrometer scale, as well as the precise comparison of different phenotypes or species. To date, we only show single-view, i.e. incomplete, reconstructions of this moth as proof-of-concept. Nevertheless, these single view measurements were repeated for different distinct positions of the moth (Fig. 1 and 2), already suggesting the possibility for a completed 360°-3D-reconstruction. 13 of such created point clouds - including the two shown here - can be found in the online repository^18^ for reference and further use.

Given the rather short field-of-depth of the system, the repositioning of the tiny specimen and subsequent point-cloud alignment is extremely delicate and will be the subject of future work. In turn, this would open up quantitative volumetric measurements of different specimen, e.g. from different habitat zones with morphological adaptations or other phenotypic classes^5^. Additionally, further improvements in the structured illumination approach could enhance the resolution towards even finer details. The presented custom calibration successfully corrected issues encountered with standard libraries, enabling accurate determination of the extrinsic parameters. Our future work will focus on optimizing the system by adapting the setup for a robust 360°-reconstruction from consecutive single-view 3D point clouds. The compact and straightforward nature of the current setup could also enable its field deployment and thus extending the approach to other biological specimens.

### Reconstruction of reference objects of known dimensions

With this framework different objects, such as a commonly used reference plane with a surface roughness *<* 1 *μm*, a microscope alignment plate with a millimeter scaling 3 (a) and a 3D-printed resin sphere could be properly reconstructed (see Fig. 7 (b) and (d)).

We used the point cloud of the alignment plate (Fig. 3 (a), Fig. 5 (b)) to calculate the scaling of the system in real world coordinates, resulting in a rescaling factor of 1.9928. The system’s performance was validated using the calibrated reference plane (Fig. 3 (b)), achieving a standard deviation of 4.46 *μm* for the full field of view while only removing outliers with a correlation value^7,8,11,20^ below 0.9. All reconstructions can be found in the supplementary file^18^.

**Figure 3.**
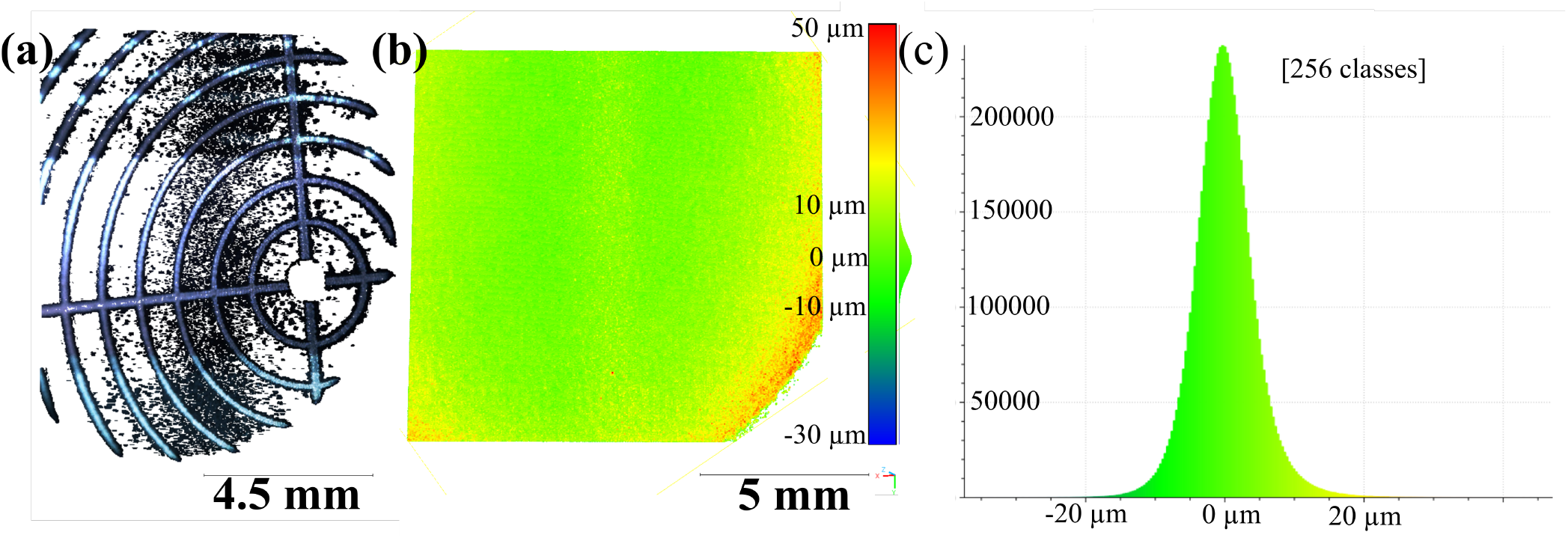
(a) 3D-reconstruction of lens mount alignment plate (see Fig. 5, Thorlabs, LMR1AP, 1 mm spacing between circles) used to estimate scale. Please note that the goal of the reconstruction was to find the bright lines not the strongly absorbing dark background. We were able to reconstruct some of those points as well, showcasing well-aligned reconstruction results even for uncooperative surfaces. (b) 3D-reconstruction of reference plane with a roughness *<* 1 *μm*. Here the entire field of view has been shown in a color scale, that represents the distance of each point from the best fit plane. The deviation increases slightly to the borders of the measurement field, as known from photogrammetry. The missing part in the lower left can be explained with a slight miss alignment of illumination and field of view. (c) Histogram of the plane reconstruction, showcasing how the standard deviation is formed from these results. Point clouds are available in the supplementary file^18^.

## Discussion

In conclusion, we have set up and adapted a telecentric stereo 3D measurement system that offers a unique combination of high resolution and large measurement volume when applied to biological photogrammetry, enabling precise volumetric analysis of specimens. Potential sources of error include subpixel inaccuracies in correspondence detection, 3D printing deviations of the reference sphere, and the delicate positioning of biological specimens. Future improvements will include automated alignment procedures and statistical analysis of volumetric precision to further quantify the measurement uncertainty. Compared to conventional stereophotogrammetric setups, our telecentric system already achieved an isotropic single digit micrometer resolution, i.e. in principle in the sub-cellular range, without additional post-processing. This is in good agreement with state-of-the-art telecentric stereo systems, but its use for biological specimens is novel. Our method, validated through the 3D reconstruction of reference objects and specimen, demonstrates the potential for applications in developmental biology and other fields, where accurate, quantitative 3D data is essential. This work paves the way for future advancements in combining macroscopic and microscopic imaging techniques, offering a new perspective on the study of morphological phenotypes in various species and potentially a plethora of adjunct fields of study.

## Methods

### Workflow Summary

The methodological workflow for the telecentric stereo 3D measurement process is illustrated in Figure 4. It consists of the following main steps:

**Figure 4.**
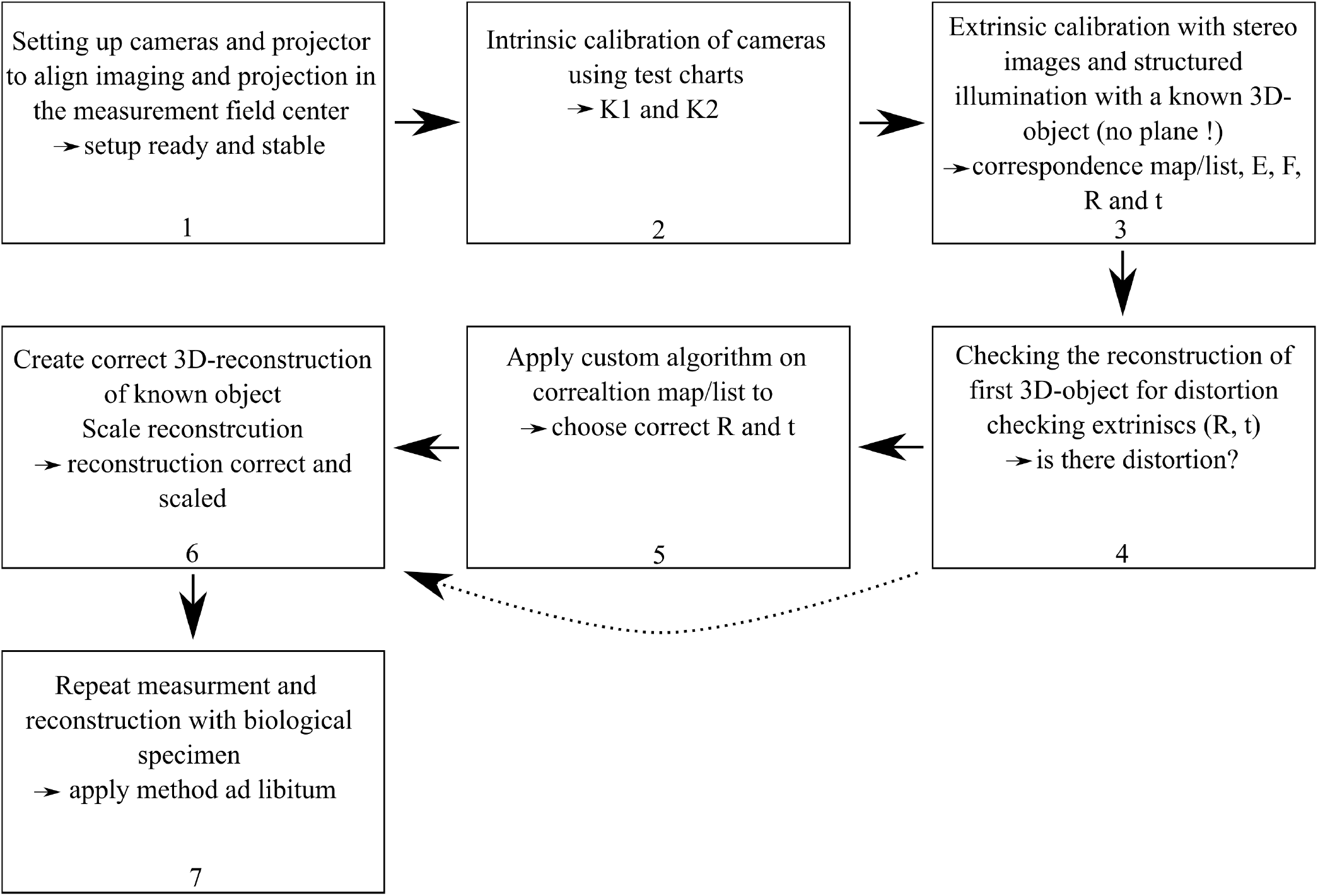
Flowchart representing the steps necessary to build and if needed correct the 3D-reconstruction methods applied here. Please note that step 5 can be skipped if the reconstruction is undistorted after step 4.

1. **System setup:** The cameras and structured light projector are mechanically aligned such that the projection axes and the optical axes of both telecentric lenses intersect at the center of the intended measurement volume. This ensures a mechanically stable and well-defined overlap region. Needed time: 1 to 4 hours, depending on user experience (DOUE).
2. **Intrinsic calibration:** Using microscopic test charts and grid targets, the intrinsic parameters of both cameras (matrices *K*_1_ and *K*_2_) are estimated to account for magnification and lateral resolution. Needed time: 1 to 4 hours (DOUE).
3. **Extrinsic calibration – initial step:** Stereo image sequences of a known 3D reference object (e.g., a resin-printed sphere) under structured illumination are captured. Based on subpixel-accurate correlation, an initial set of extrinsic parameters (rotation *R* and translation *t*) is derived using standard procedures (e.g., from the essential and fundamental matrix). Needed time: 5 to 10 minutes measurement time, depending on algorithm seconds^20^ to 30 minutes for the extrinsic calibration.
4. **Distortion check:** The reconstructed point cloud is compared against the known geometry of the reference object. Significant deviations or warping indicate a model mismatch caused by standard pinhole assumptions. Needed time: 5 to 10 minutes.
5. **Model correction:** A custom algorithm is applied to resolve the ambiguity introduced by mismatched assumptions in the extrinsic calibration. With the help of the algorithm (and this manuscript) the user can identify the correct combination of *R* and *t* that matches the telecentric projection geometry. 2 Minutes to 30 minutes (DOUE).
6. **Final 3D reconstruction and scaling:** The corrected extrinsic parameters are used to generate an accurate, undistorted point cloud. Scaling is performed using features with known real-world dimensions, enabling quantitative interpretation. Needed time: 30 seconds for each reconstruction.
7. **Application to biological specimens:** Once validated, the system is applied to fragile biological samples. Multiple measurements may be repeated at different orientations to allow for full 3D reconstruction or volumetric analysis. Depending on sample properties and experience level of user 5 to 20 minutes per sample and reconstruction.

### Setting up a telecentric stereo system with structured illumination I - Optical system setup

Photogrammetric 3D-measurement techniques with structured illumination are state of the art for a lot of applications. Also the more specialized sub-methodology - using telecentric cameras instead of ‘regular’ or endocentric cameras - has been applied successfully also with structured illumination. However, these instances have been performed by experts of image processing and optical metrology. Researchers from other fields with limited experience in optical metrology may benefit from a simplified, pre-configured system. Here, we will try to explain and to guide through the steps necessary to setup and reproduce our experiment and point out problems, that we ran into.

The first part is assembling the mechanical setup: Our advice is, to start with the projector and one camera. Here Allied Vision (Stadtroda, Germany) Alvium cameras (1800 U-811c color C-Mount), with 2848×2848 pixels each and a pixel pitch of 2.74 *μm* have been used. The telecentric lenses were of the type LM1123TC from Kowa (2/3” 0.69-0.88x) and a low cost off-the-shelf projector (Technaxx Mini LED Beamer TX-113) was in use. The projector has to be connected to the same computer, the cameras are connected to. Also, at least two additional optics, for example lens doublets with 2” diameter and a focal length of 75 *mm* (in this case Thorlabs AC508-075-A-ML) can be used to minimize the field of projection to the desired size of 10*mm* x 10*mm*. A blank piece of white paper can be used as a screen. One of the cameras (with a telecentric objective) can be used here to evaluate contrast, field of view and depth of field. The working distance with the system in use here was 12 *cm*. We experienced that an almost completely closed aperture on the imaging as well as on the projection size yields the best depth of field and depth of projection - but also requires significantly more imaging time of up to a few seconds per images/images pair for the biological sample. However, these measurements created the best images and also the best 3D-reconstruction. When the center of projection and the center of imaging of the first camera are overlapping, the second camera can be added to image the same center - the center of the measurement field (see Fig. 6 a). It might be of interest to remark, that the achievable 3D precision is proportional to the sine of the stereo angle of the system. A larger angle also reduces the overlap of the fields of view and increases the risk of occlusions within the measurement volume. Therefore a sweet spot for each application has to be found. Here we realized an angle of approximately 38.5° and a baseline length of approximately 9 *cm*. After the physical setup the system has to be calibrated, namely intrinsically, so in terms of imaging process by each camera, and extrinsically, in terms of the position of the cameras towards each other.

### Setting up a telecentric stereo system with structured illumination II - System calibration

The imaging process by common or endocentric lenses in first order approximation can be mathematically described with the pinhole model, yielding in a 3 × 3 matrix (*K*_*Endo*_) with the focal length *f* in the spatial directions *x* and *y*, as well as the projected position of the pinhole in the camera plane with the coordinates (*m*_*x*_,*m*_*y*_) (see Eq. 1). Also, a shearing parameter *s* can be added to accommodate an angle between the pixel orientation in the lateral spatial directions. In comparison, the imaging process with a telecentric lens, as used in this study, can be described with a matrix *K*_*Tele*_ that has nonzero values only on the main diagonal (see Eq. 2). These are the magnification divided by the pixel pitch in *x* and *y* and a 1^14–16^.

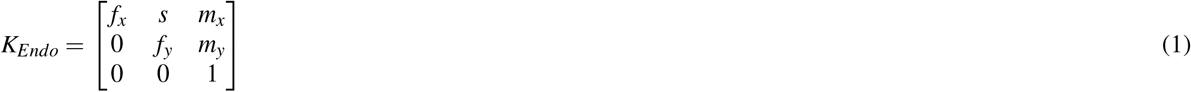

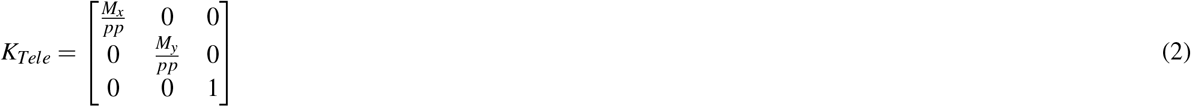

This difference mathematically resembles the idea, that the viewing rays of endocentric cameras are meeting in exactly one point in space, the pinhole. On the other hand telecentric imaging resembles a parallel projection, so that the meeting point of the parallel viewing rays lies at infinity. Like in all 3D-measurement approaches, precise calibration of the system, i.e. the determination of *K*, results in high accuracy and precision, crucial for reconstructing delicate structures like moth bodies. Especially in the presented case, i.e. the rededication of a known method and open-source libraries to a customized telecentric imaging system, the calibration is crucial.

The first part of calibration in photogrammetry is finding an appropriate imaging matrix according to Eq. 1 (right) for each camera. To intrinsically calibrate, marked reference planes (graph paper and checker boards as well as a microscopic test chart, see Fig. 5) have been used to identify the magnification and the lateral resolution by comparing real size of objects to imaged size of objects^14–16^ (see supplementary file^18^). For non-expert users a simplified version of finding the magnification could be using graph paper and taking images of it when orientated perpendicular to the viewing direction of one of the cameras and then repeating this for the second camera. With the knowledge of the pixel pitch a comparison of the length of the grid marks in pixels (multiplied with the pixel pitch) can be compared with the real size and such found an approximation for the magnification in *x* and *y*. Please note that this simplified method might be imprecise but was found to deliver comparable results to more sophisticated approaches by the authors of this manuscript.

**Figure 5.**
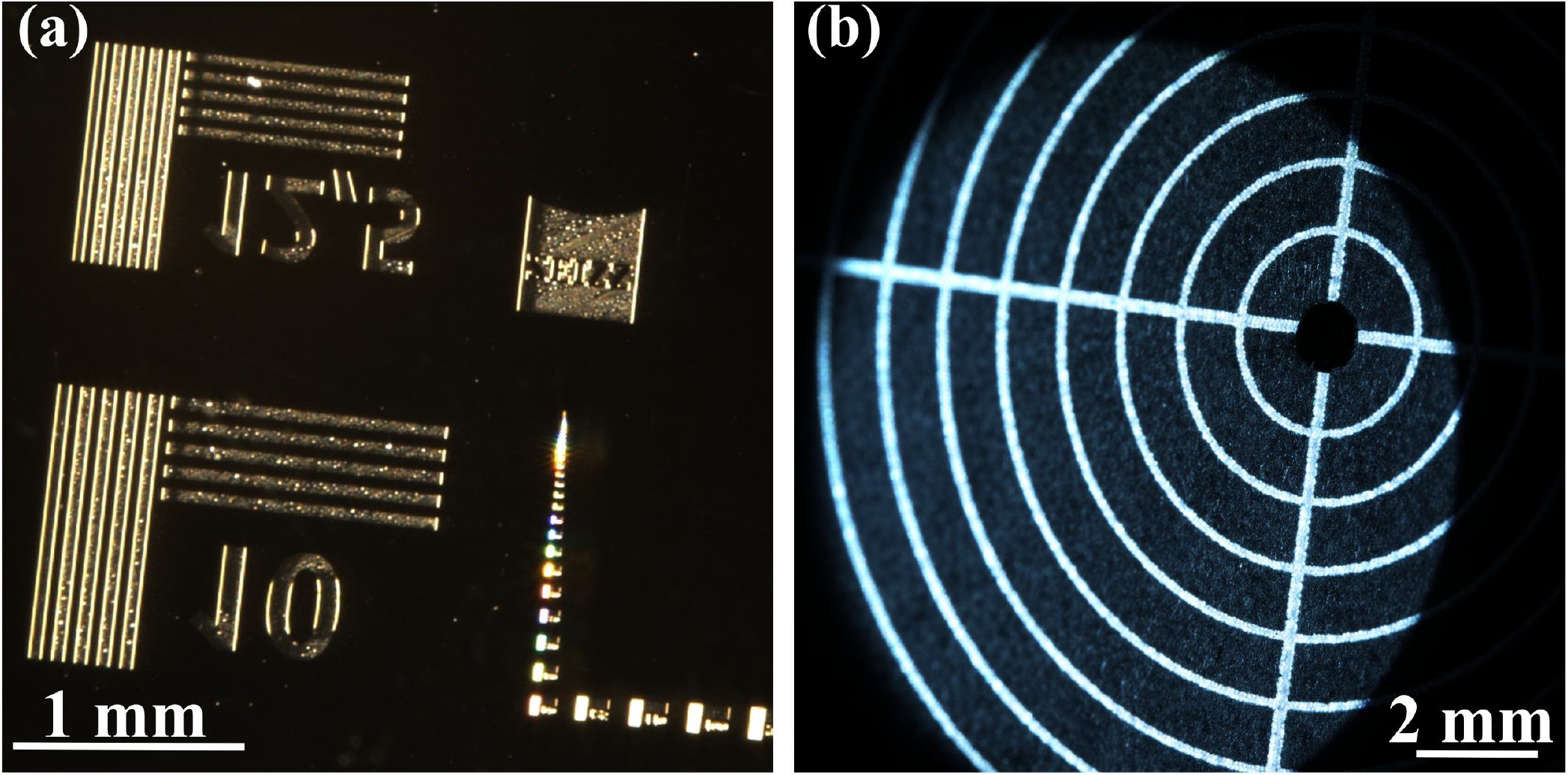
(a) Image (cropped) of microscopic reference chart used to estimate the lateral resolution and magnification. (b) Images of a lens mount alignment plate (Thorlabs, LMR1AP, 1 mm spacing between circles, left) used to estimate scale. The complete images are available in the supplementary file^18^.

The pixel pitch has been taken from the camera manufacturer. We found the effective magnification of the lenses to be 1.46 and thus the lateral resolution to be 8 *μm* (using 5 (a)). Here, we took the size of the reference structure in the images in pixel into account and evaluated the resolution from the visibility of the smallest resolved structure. The intrinsic matrices *K*_1_ and *K*_2_ for both cameras were found and are presented in Eq. 3 and Eq. 4.

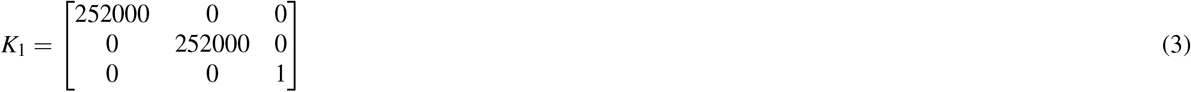

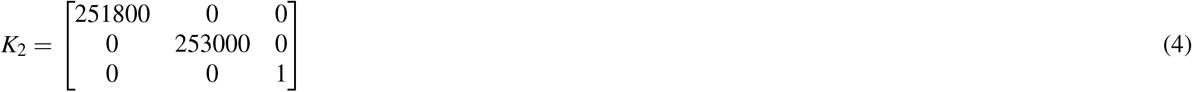

The alignment plate (Fig. 5 (b)) on the other hand, was later used to retrieve the real world scale from the setup. Please note that this was done after the 3D-reconstruction of the alignment plate, also to support the reliability of the system.

The straightforward 3D measurement setup (see Fig. 6 (a)) is comprised of the two mentioned cameras and a commercial projector with a magnifying lens assembly to fill a 11 *mm* × 11 *mm* × 6 *mm* measurement volume. The second step is to find the extrinsic parameters (extrinsics), i.e. the rotation and translation from the first image plane (camera 1) to the second one (camera 2). Here, a measurement of a resin printed sphere with a radius of 2.5 *mm* has been taken with 100 stereo images and varying structured illumination, comparable to already published studies^7,8,11^, see Fig. 6 (b), (c). A recent, openly accessible reconstruction algorithm can be found here^20^. This rapidly fast algorithm is difficult to understand and implement for non-expert users.

**Figure 6.**
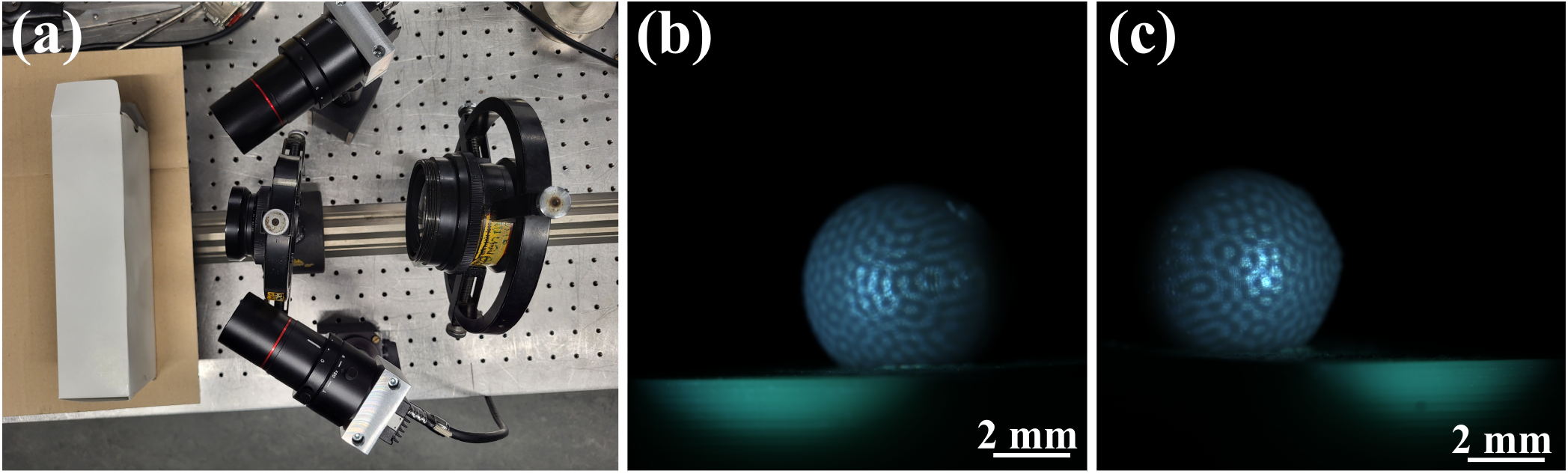
Measurement setup (a), with two cameras with telecentric lenses and two projecting lenses (projector outside of image). (b) 3D-printed sphere (radius = 2.5 *mm*) as imaged from first and (c) second camera under structured illumination (supplementary file^18^).

### Correcting the distortion and scaling

Using the temporal correlation search for correspondence pairs^7,8,11^ yielded sub-pixel accurate results. However, using these results to calculate the initial extrinsic parameters of the measurement system resulted in heavily distorted 3D reconstructions when compared to a best fit sphere (see Fig. 7 (a) and (c)).

**Figure 7.**
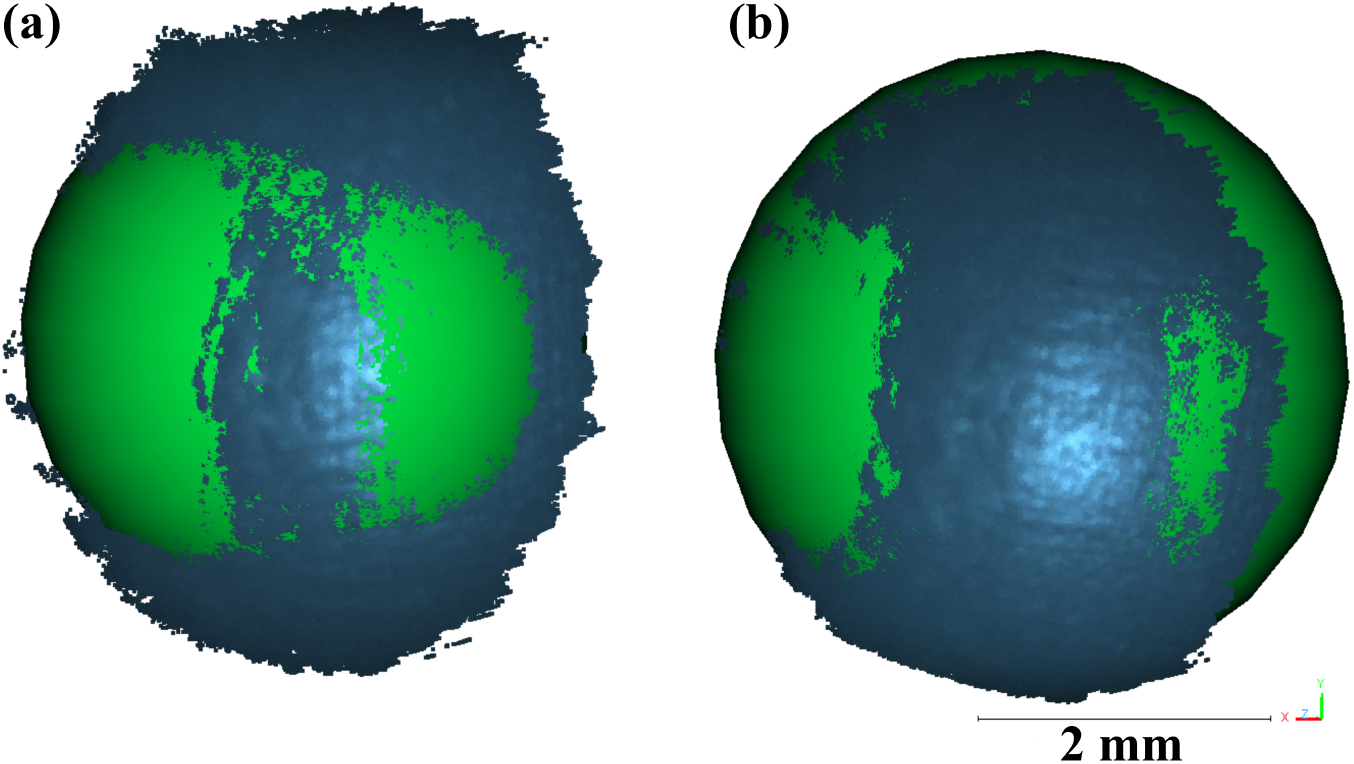
(a) Distorted and unscaled point cloud/3D-reconstruction of the resin sphere in original, bluish colors - see Fig. 6 (b) and (c) - and a best-fit sphere (green). (b) Undistorted and scaled 3 point cloud/3D-reconstruction of the resin sphere in original, bluish colors and a best-fit sphere (green). The data was rendered in CloudCompare^19^. Points from the holding platform (Fig. 6 (a) and (b)) have been removed. *x, y* and *z* -coordinates shown in tripod lower right corner. Point clouds are available in supplementary file^18^.

The distortion in this case were obvious due to the vast misalignment of the reconstruction with a best fit sphere. We also encountered distorted reconstruction when putting other samples under test however. In this context we present only the sphere, as a clear reference for a misreconstruction model. Non-expert users can however take other objects of known (relative) dimensions to check for a correct reconstruction. In some cases, the standard calibration (step 4) may already yield the correct extrinsic parameters. In such situations, step 5 can be skipped, as no distortion is observed in the initial 3D reconstruction. The significant distortion encountered by us is not surprising, as standard libraries that we aim to utilize to enable open-source access and low entry barriers for non-experts such as life-scientists, e.g. OpenCV, presume non-telecentric/endocentric imaging systems that can be described by a pinhole model. Thus, they prove to be inadequate if applied naively on the extrinsic calibration, leading to distorted reconstructions (Fig. 7 (a)). In mathematical terms, a set of two rotational matrices and two translation vectors is found when calibrating extrinsically this way. However, due to the inherent model mismatch, the algorithm will produce likewise mismatched extrinsics by discriminating the wrong set of matrix and vector. To address this, we rewrote a custom algorithm inspired by standard literature of the field^21^ that allowed to correctly choose from the possible set of extrinsics to accommodate the model mismatch (compare Eq. 1, Eq. 2 and see the supplementary file^18^). Utilizing the correct extrinsic parameters consistent with the telecentric model, the identified pixel-pair correspondences were then straightforwardly used to reconstruct undistorted point clouds as in^7,8,11^ via triangulation. Please note that the two possible translation vectors only differ by sign while the two options for the rotational matrix represent a real second perspective and the reprojection in the first camera view. In principle it is not important whether ^*′*^*t*^*′*^ or ^*′*^*− t*^*′*^ are chosen. This creates reconstructions with a sign change on the *x* and *z* axes but geometrically correct. Choosing the wrong rotational matrix on the other hand leads to the described distortion. As a rule of thumb (for non-expert users) - the rotational matrix that shows greater deviation from the identity matrix is (usually) correct.

The sphere was reconstructed using a correlation threshold^7,8^ of 0.3 and removing points from the holding platform. For a fixed radius of 2.5 *mm* a standard deviation from sphere form of 63.1 *μm* was found. Note that only a sector of the resin sphere was visible within the overlapping field of view, limiting a fit with a flexible radius to a portion of the full sphere. The actual diameter of the sphere may differ slightly from 5 *mm*, in addition.

## Acknowledgements

This work has been supported by the German Federal Ministry for Economic Affairs and Climate Action (BMWK) within the Promotion of Joint Industrial Research Program (IGF) as part of the research project (IGF 22462 BR) by the Association F.O.M., as well as the German ministry of Education and Research (BMBF) within the Research Program Quantum Systems under the project number 13N16890.

The authors would like to thank Dr. Peter Kühmstedt for his insights to telecentric measurement and to Dr. Daniel Girardeau- Montaut and colleagues for providing CloudComapre^19^.

## Author contributions statement

A.W.S. and C.F. conceived the experiments, A.W.S. and A.O. conducted the experiments, A.W.S analyzed the results. G.J.G. and M.G. wrote and provided programs for image capturing and 3D-reconstruction. D.G. and G.B. provided probes and expertise into lepidopetrans. All authors reviewed the manuscript.

## Additional information

**Accession codes** Data underlying the results presented in this paper are available in a supplementary file - separated in 3 sub-files 1, 2 and 3. Here supplementary file 1 includes the images shown in the manuscript, file 2 includes the 3D point clouds and file 3 the program and matrices in use to calculate the extrinsic matrices from a set of corresponding points and one to estimate a best fit sphere. A correspondence search algorithm can be found in^20^ or provided by the authors upon reasonable request.

## References

1. Bresee, C. S., Belli, H. M., Luo, Y. & Hartmann, M. J. Comparative morphology of the whiskers and faces of mice (mus musculus) and rats (rattus norvegicus). J. Exp. Biol. 226 (2023).

2. Weiler, S. et al. A primary sensory cortical interareal feedforward inhibitory circuit for tacto-visual integration. Nat. Commun. 15, 3081 (2024).

3. Ströbel, B., Schmelzle, S., Blüthgen, N. & Heethoff, M. An automated device for the digitization and 3d modelling of insects, combining extended-depth-of-field and all-side multi-view imaging. ZooKeys 1 (2018).

4. Ancajima, G. P., Eloi, I. & Duarte, M. Sexual dimorphism and allometric patterns in hawkmoth epiphyses (lepidoptera: Sphingidae). Sci. Reports 15, 11405 (2025).

5. Brehm, G., Zeuss, D. & Colwell, R. K. Moth body size increases with elevation along a complete tropical elevational gradient for two hyperdiverse clades. Ecography 42, 632–642 (2019).

6. Mungee, M. & Athreya, R. Rapid photogrammetry of morphological traits of free-ranging moths. Ecol. Entomol. 45, 911–923 (2020).

7. Harendt, B., Große, M., Schaffer, M. & Kowarschik, R. 3d shape measurement of static and moving objects with adaptive spatiotemporal correlation. Appl. Opt. 53, 7507–7515 (2014).

8. Stark, A., Wong, E., Babovsky, H. & Kowarschik, R. Subjective speckle suppression for 3d measurement using one-dimensional numerical filtering. Appl. Opt. 58, 9473–9483 (2019).

9. De Leo, N., Chimenti, C., Maiorano, L. & Tamagnini, D. Protocol for 3d photogrammetry and morphological digitization of complex skulls. STAR protocols 6, 103572 (2025).

10. Beri, A. et al. Quantitate evaluation of photogrammetry with ct scanning for orbital defect. Sci. Reports 14, 3104 (2024).

11. Heist, S. et al. Gobo projection for 3d measurements at highest frame rates: a performance analysis. Light. Sci. & Appl. 7, 71 (2018).

12. Hu, Y. et al. Structured light 3d imaging instrument for biological tissues with potential application in telemedicine. IEEE Transactions on Instrumentation Meas. 73, 1–11 (2023).

13. Bolaños, L. A. et al. A three-dimensional virtual mouse generates synthetic training data for behavioral analysis. Nat. methods 18, 378–381 (2021).

14. Zhang, S., Li, B., Ren, F. & Dong, R. High-precision measurement of binocular telecentric vision system with novel calibration and matching methods. ieee access 7, 54682–54692 (2019).

15. Hu, Y. et al. Calibration and rectification of bi-telecentric lenses in scheimpflug condition. Opt. lasers engineering 149, 106793 (2022).

16. Liu, H. et al. Epipolar rectification method for a stereovision system with telecentric cameras. Opt. Lasers Eng. 83, 99–105 (2016).

17. Chen, C., Kong, Y., Wang, H. & Zhang, Z. A high-accuracy calibration method for a telecentric structured light system. Sensors 22, 6370 (2022).

18. Stark, A. W. Telecentric reconstructions and data. figshare, 10.6084/m9.figshare.c.7652423.v1 (2025).

19. Girardeau-Montaut, D. Cloudcompare 3d point cloud and mesh processing software open source project (2025). Accessed: 2025-03-18.

20. Liebender, C., Bleier, M. & Nüchter, A. libbicos–an open source gpu-accelerated library implementing binary correspondence search for 3d reconstruction. The Int. Arch. Photogramm. Remote. Sens. Spatial Inf. Sci. 48, 57–64 (2024).

21. Hartley, R. & Zisserman, A. Multiple view geometry in computer vision (Cambridge university press, 2003).

